# Moving Heads, Moving Sounds: Systematic Errors in Auditory Motion Perception

**DOI:** 10.64898/2026.11.05.764571

**Authors:** Timo Oess, Laurence R. Harris, Marc O. Ernst

## Abstract

The movement of moving objects seen during self-motion is incorrectly perceived as a result of errors in parsing the visual motion arising from the retina and from information about the movement of the head and eyes in space^1^. The perception of motion of auditory targets may be even more challenging as previous research suggests that the auditory system lacks dedicated low-level speed processing mechanisms. This study investigates how the perception of a moving auditory sound source is influenced by concurrent head movements, drawing comparisons with established visual motion perception findings and illusions, such as the Filehne Illusion. Participants were asked to move their heads at a fixed speed, either horizontally (left to right or right to left) or vertically (up to down or down to up), while indicating the perceived direction of a moving sound source by using a joystick. A physical sound source was moved silently at a constant speed in fronto-parallel 2D space using a 3 × 3×3m cable robot. Results showed that head movements created a perceptual bias of the perceived direction of the moving sound source in the opposite direction to the head movement. Responses were less precise during horizontal than vertical head movement, whereas overall directional accuracy did not differ reliably between movement planes. The magnitude of the directional error also depended on the direction of head movement, consistent with the different cues available for horizontal and vertical localization (binaural vs. spectral cues). These findings suggest that similarly to how visual motion perception is distorted by extraretinal signals and the movement of the eyes, so too is auditory motion perception influenced by movement of the ears, revealing errors in the integration of proprioceptive, vestibular and auditory cues that distort the perceived direction of a moving sound source.

## Introduction

Imagine walking along a quiet country road when a bicycle approaches from behind. To check whether it will pass you on the right or left side, you turn your head expecting your percept of the bike’s location in space to remain accurate. In order to do this, the auditory system must disentangle the auditory motion cues originating from the bike from those generated by the listener’s own head movements (see Fig. X). That is, the brain needs to translate the perceived signals from a head-centered reference frame, in which the bike seems to move opposite to the head movement, to a world-centered reference frame, in which the head movement is accounted for and the bike‘s actual movement is perceived correctly.

**Figure X:**
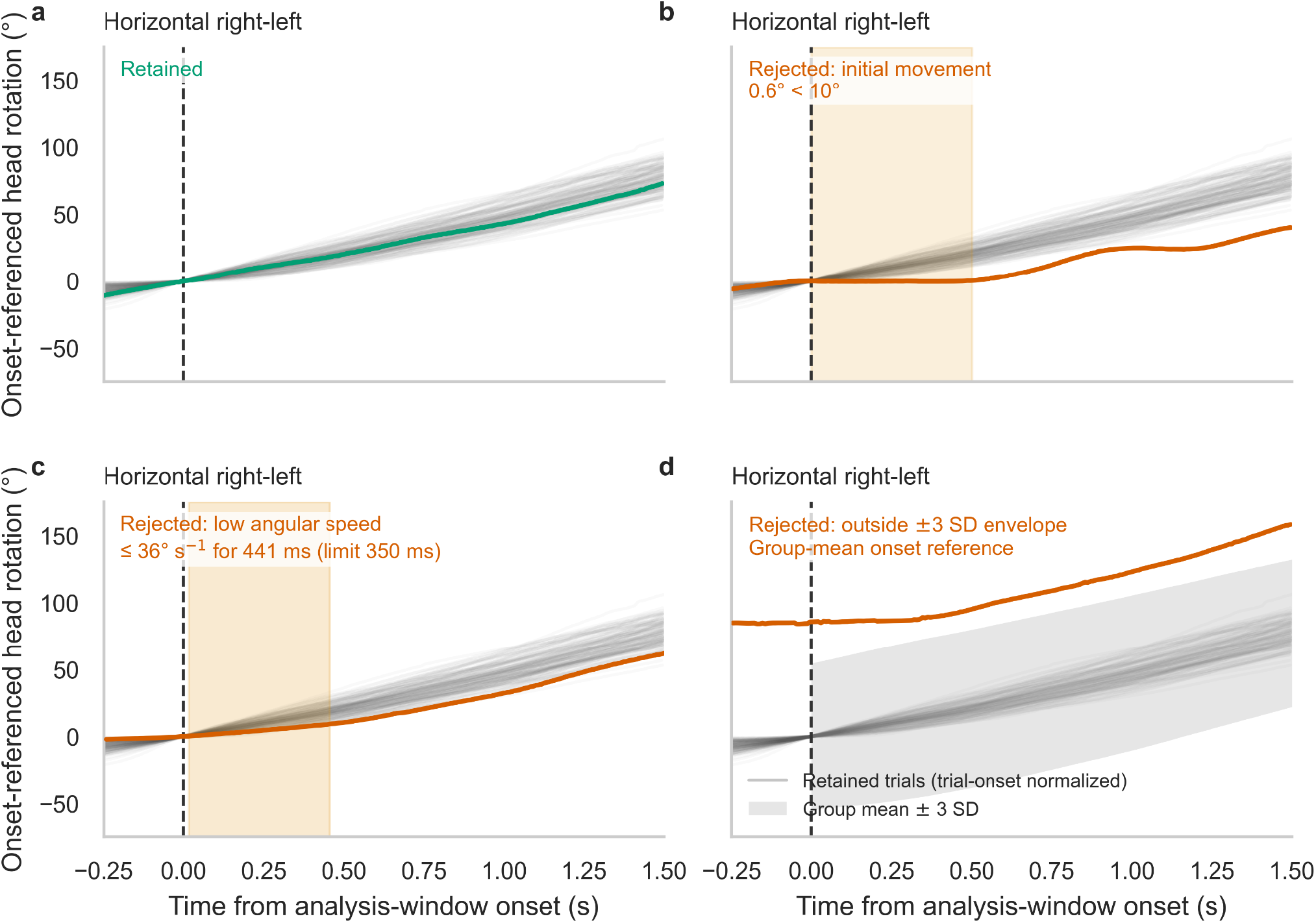
Representative head-rotation trajectories and trial-exclusion criteria. Head rotation is shown from 250 ms before to 1.5 s after the onset of the analysis window (vertical dashed line).

This problem of motion parsing—distinguishing self-induced from external motion—has been well described in vision, where failures lead to perceptual biases such as the Filehne Illusion^2^. In audition however, the mechanisms underlying motion perception are less well understood, particularly because spatial cues about an auditory object are not yet present at the level of the receptor (compared to the retina) but need to be computed at a later processing stage^3,4^. Additionally, there are different cues for localizing, and thus determining the motion direction of an auditory object, in the horizontal plane and in the vertical plane. In the horizontal plane, computed binaural cues, based on the difference of arrival time and/or intensity between the left and right ear, interaural time differences (ITD) and interaural level differences (ILD), are used^5,6^. However, these cues can be ambiguous (see “cone of confusion”^7^) and remain constant for sound sources on the mid-sagittal plane. Thus, the auditory system additionally takes advantage of the filtering properties of the pinna to determine the elevation of a sound source and resolve front-back confusions^6,7^. Head movements themselves provide a further, dynamic cue that can help resolve such ambiguities for stationary sound sources^8,9^. These factors render the underlying processes of auditory motion perception during self-motion different from vision. It is therefore unclear whether the auditory system is able to compensate for self-motion as well as the visual system can. In this study, we set out to determine the compensation ability of the brain for moving sound sources under self-motion.

Freeman and colleagues^10^ demonstrated that during horizontal head movements, listeners exhibit systematic biases in judging the motion of sound sources which can be quantified by a “compensation gain”. Their study was restricted to the horizontal plane and to conditions in which the directions of head and sound source motions were aligned. As such, the effects they observed primarily reflected distortions in binaural cues—interaural time differences (ITD) and interaural level differences (ILD).

It remains unclear how the auditory system compensates when sound sources move in more naturalistic trajectories, extending beyond the horizontal plane or involving vertical head movements. Under these conditions, the brain must rely not only on binaural information but also on spectral cues, and on dynamic combinations of both.

In the present study, we present real physical motion of a loudspeaker using a cable robot in three-dimensional space (Suppl. Fig. X). This setup allowed us to test auditory motion perception across 16 distinct directions, including both concurrent and orthogonal head– source motion combinations (see Figure 2B). A central question was whether compensation mechanisms generalize to vertical head movements, where ITD and ILD cues—the most reliable spatial indicators—remain relatively undisturbed. We hypothesized that head motion would bias perceived source motion across all directions, but that perceptual accuracy would be greater during vertical head movements due to more stable interaural cues.

**Figure 1:**
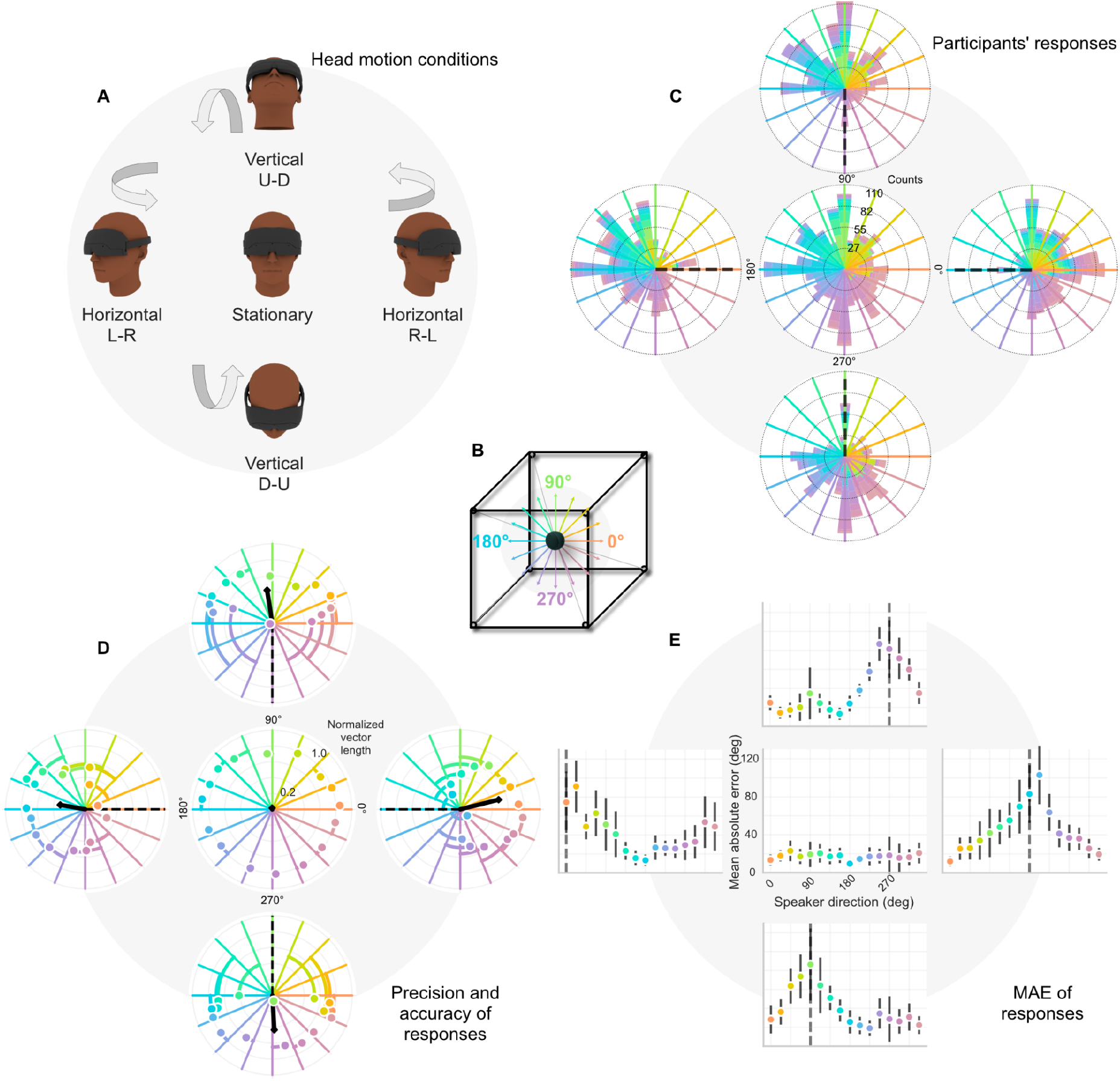
Experimental design and perceived sound-source localisation accuracy across head-motion conditions. (A) Head motion conditions. Participants were blindfolded and wore a head-mounted motion-tracking device. On each trial, they were instructed to perform a prescribed continuous head movement — independently of the sound — according to one of four head- motion conditions: Horizontal Left-to-Right (Hori L-R), Horizontal Right-to-Left (Hori R-L), Vertical Up-to-Down (Vert U-D), or Vertical Down-to-Up (Vert D-U). In the Stationary control condition, no head movement was performed. Simultaneously, a real sound source moved along a randomly selected direction, and participants indicated their perceived direction of sound-source motion by pointing a joystick at the end of each trial. The results are arranged to match the same spatial layout. (B) Spatial reference frame. The sound source moved along one of 16 evenly spaced angular directions (22.5° increments) within the horizontal or vertical plane, selected randomly on each trial. Directions are defined as: 0° = right, 90° =up, 180° = left, 270° =down. (C) Participants’ responses. Stacked polar histograms showing the distribution of reported perceived sound-source directions across all trials and participants, separately for each condition. Each spoke corresponds to one of the 16 possible speaker directions; bar height encodes response count, with stacked segments representing individual trials. Colours encode speaker direction (as shown in **B**). A perfectly accurate response pattern would yield responses uniformly distributed and aligned with their respective target direction spokes (colors perfectly matching). (D) Precision and accuracy of responses. Vector-sum polar plots for each condition. For each of the 16 speaker directions, the vector sum of responses across participants was computed; the direction of the resultant vector reflects the mean perceived sound direction and its normalised length (0–1) reflects response precision across trials and participants. Coloured dots show per-sound motion direction vector-sum endpoints; the black arrow shows the group-level mean vector. The dashed line marks the prescribed head-motion axis for each condition. Annotated values indicate the normalised vector-sum magnitude for selected directions. (E) Mean absolute error (MAE) of responses. Mean absolute angular deviation between the reported perceived direction and the actual speaker direction (°), plotted as a function of speaker direction (0°–360°), separately for each condition. Each data point represents the group mean; error bars indicate 95% confidence intervals (1.96 × SE across participants). The y-axis is bounded at 120°. The vertical dashed line marks the prescribed head-motion axis for each condition.

**Figure 2:**
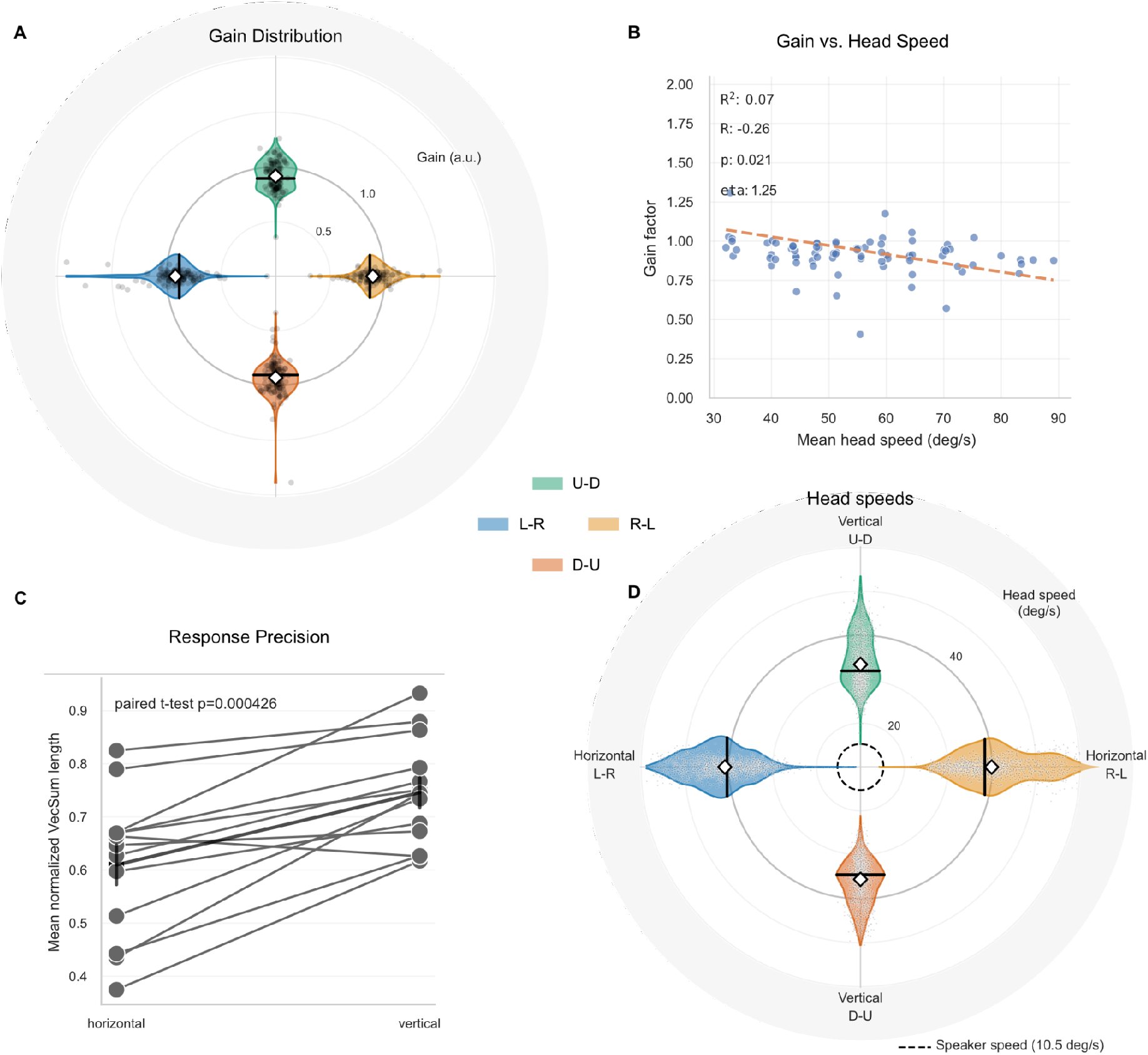
Head movement speed and gain factor across movement conditions. (A) Mean angular head speed (deg/s) during each movement condition (Hori L-R, Hori R-L, Vert U-D, Vert D-U), shown as violin plots with individual participant data points overlaid. The dashed horizontal line indicates the speaker movement speed (10.5 deg/s). Participants moved their heads considerably faster than the sound source across all conditions. (B) Distribution of gain factors per movement condition shown as violin plots, with individual trial values overlaid. The gain factor is the fitted scaling of the residual head-motion contribution in the vector model V(g) = S + (g − 1)H, where g = 1 denotes full compensation (veridical perception of source motion) and g = 0 denotes no compensation. Mean and median values are annotated for each condition. The dashed horizontal line marks unity gain (gain = 1). (C) Gain factor as a function of mean head speed (deg/s), pooled across all movement conditions and restricted to speaker directions perpendicular to the head motion axis (0°/180° for vertical, 90°/270° for horizontal conditions). Each data point represents the fitted gain estimate for a specific combination of movement condition, speaker direction, and head-speed bin. The dashed line shows the overall linear regression fit. A significant negative relationship was observed (R^2^ = 0.07, R = −0.28, p = 0.021), indicating that faster head movements were associated with lower gain. (D) Comparison of mean normalized vector sum length (precision) between horizontal (Hori L-R and Hori R-L combined) and vertical (Vert U-D and Vert D-U combined) conditions. Individual participants are connected across conditions. A paired t-test revealed a significant difference between orientation axes (t(12) = 3.93, p = .002, dz = 1.09), with vertical conditions yielding higher precision than horizontal conditions.

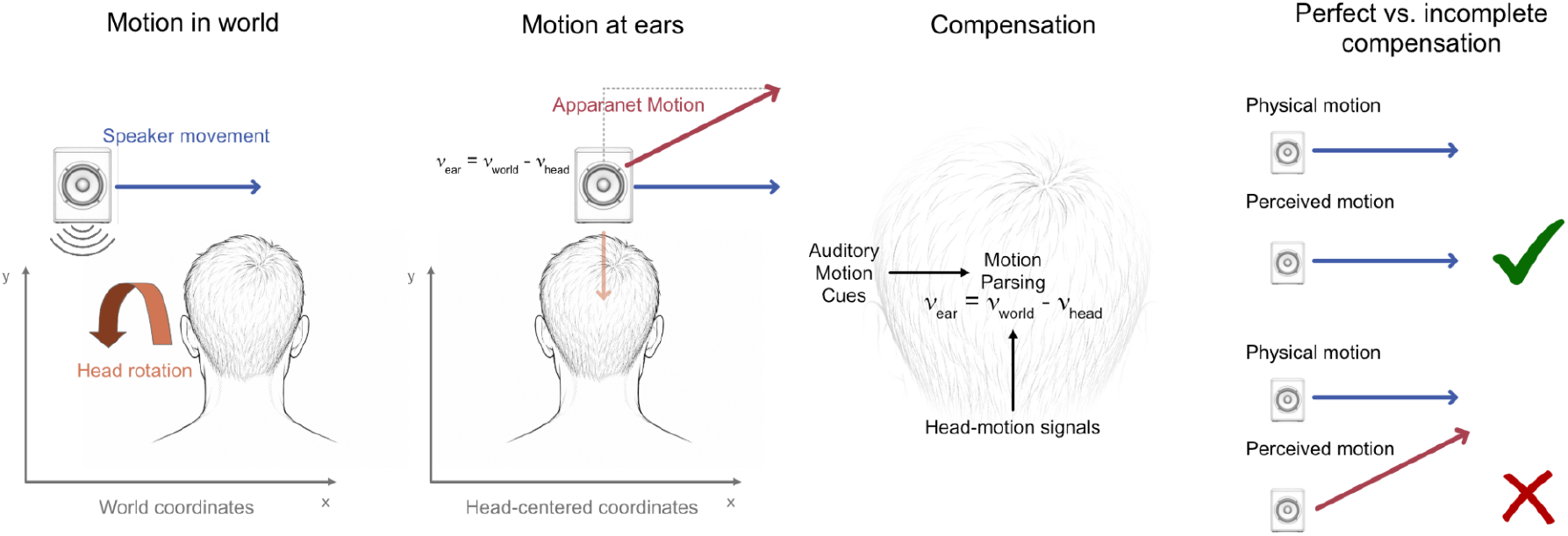

Figure: Transforming perception into world-centered coordinates

## Results

To investigate the influence of head movements on the perception of moving auditory objects, we tested participants in an auditory motion-direction perception task. Participants, blindfolded throughout the experiment, were seated in front of the 3D motion setup (Fig. 1 B) and asked to indicate the perceived direction of an auditory stimulus after each trial. We defined 16 predefined two-dimensional linear trajectories (see different colors Fig. 1B), and in each trial the speaker moved along a randomly chosen trajectory. The stimulus consisted of 1.5 s of continuous white noise with onset and offset ramps, played shortly after the onset of speaker movement. We tested five conditions in total, in which participants either kept their heads stationary (S) or moved them from left to right (L-R), right to left (R-L), up to down (U-D), or down to up (D-U) (Fig. 1A). The responses of all participants were combined for analysis (see the supplementary materials for single-participant performance).

### Participants’ responses are biased by head movements

Based on previous research on visual^1^ and auditory^4,10^ motion tracking during self-motion, we hypothesized that participants’ responses would be systematically biased by head movements. To assess potential response biases, responses were binned (bin size = 10°) and visualized using color-coded polar histograms (one per condition), in which each loudspeaker motion direction was assigned a unique color and corresponding response bins were colored accordingly (Fig. 1C).

Performance in the baseline (stationary head) condition was high, indicating that, in the absence of self-motion, participants were able to reliably report the direction of a moving sound source. In contrast, the histograms for head-motion conditions reveal a pronounced clustering of responses in the direction opposite to the actual head movement (indicated by the dashed line), with a near absence of responses in the direction of head movement. This distribution provides strong evidence that head motion systematically biases the perceived direction of sound source movement.

This effect is further reflected in the mean absolute response error (defined as the angular difference between participant response direction and speaker movement direction). Errors are largest for speaker motion directions that are aligned with, or close to, the direction of head movement (Fig. 1E), while remaining minimal in the baseline condition.

Interestingly, for vertical head movements (U-D and D-U conditions), response errors also show a modest increase for speaker directions opposite to the head movement. This may reflect reduced reliability of spectral cues (e.g., notches and troughs) for vertical motion perception.

To quantify perceptual bias in each head-motion condition, we computed the vector sum of all responses, extracting both its direction and normalized magnitude (Fig. 1D, black line). The direction of this vector provides an estimate of the mean bias, while its magnitude reflects response precision (with a value of 1 indicating perfect precision). Across all conditions, the resulting vectors are oriented almost exactly opposite to the direction of head movement (dashed black line), further demonstrating a strong influence of head motion. The vector magnitudes indicate a reduction in precision under head-motion conditions relative to baseline.

To further examine direction-specific biases, we computed vector sums separately for each speaker motion direction (Fig. 1D, colored dots). For all conditions, these vectors are closest to the true speaker motion direction when that direction is far from the head-motion direction. This relationship is illustrated by colored arcs connecting each estimated vector to the corresponding true speaker direction; shorter arcs indicate smaller systematic errors. In addition, the distance of each vector from the origin reflects response precision, which decreases as speaker motion becomes more aligned with head movement, approaching zero for congruent motion directions.

This pattern suggests increasing difficulty in compensating for self-motion when auditory stimuli move in the same direction as the head. Consistent with this interpretation, participants reported that, in such trials, the sound source often appeared stationary and its direction of motion could not be reliably determined.

### Compensation is incomplete for all head movement directions

To quantify the extent to which participants compensated for their own head movements, we estimated compensation gain from head velocity (speed and direction), speaker velocity, and the reported response. The gain displayed here is a descriptive estimate obtained per movement condition and speaker direction from the response distribution pooled across participants; a participant-level version of this analysis, which treats the participant as the unit of inference, is reported in the Supplementary (Supplementary Fig. S3). A gain of 1 indicates perfect compensation, i.e., veridical perception of the external sound-source motion despite self-motion. Trials for which no meaningful gain could be estimated were excluded (see Methods).

The resulting gain values indicate systematically incomplete compensation across all head- movement conditions. Mean gain values were 0.87 (left-to-right), 0.93 (right-to-left), 0.86 (up- to-down), and 0.93 (down-to-up), with corresponding median values of 0.90, 0.89, 0.91, and 0.93 (Fig. 2A). A participant-level re-analysis — fitting one compensation gain per participant and condition directly to the trial-wise circular responses rather than to responses pooled across participants, and restricting estimates to geometries in which gain is identifiable — reproduced this pattern of systematic undercompensation (mean gain 0.79–0.91 across conditions; Supplementary Fig. S3).

During the training phase, participants were instructed to match their head movement speed to that of the moving sound source (∼10.5°/s). However, actual head velocities were substantially higher, averaging approximately 53°/s (Fig. 2D), with considerable variability across all conditions. This indicates that participants systematically overshot the instructed target velocity. Horizontal head movements (left-to-right: 61.8°/s; right-to-left: 59.8°/s) were, on average, faster than vertical movements (up-to-down: 46.7°/s; down-to-up: 51.1°/s).

The variability in head speeds allowed us to examine their influence on compensation performance. Trials were binned according to executed head speed (10 bins with equal sample size, n = 632 per bin), and gain values were computed separately for each head- motion condition, focusing on trials with perpendicular speaker motion (as these provide the most geometrically informative and stable estimates of gain). This binned, across-participant analysis is descriptive and exploratory — it pools trials into head-speed bins rather than treating the participant as the unit of inference — and showed a negative association between head speed and gain (Pearson’s r = −0.28, r^2^ = 0.07, p = 0.021; Fig. 2B), consistent with poorer compensation at higher head speeds. We therefore treat it as suggestive rather than confirmatory.

Another factor likely influencing compensation performance is the extent to which auditory spatial cues are confounded by self-motion. During horizontal head movements, the most reliable localization cues—interaural time differences (ITDs) and interaural level differences (ILDs)—change dynamically even for stationary sound sources. As a result, changes in these binaural cues are inherently ambiguous, reflecting a mixture of self-motion and external source motion. This ambiguity is particularly problematic when sound-source motion occurs along or near the same axis as the head movement, requiring the auditory system to disentangle two sources of similar cue variation.

In contrast, during vertical head movements, self-motion leaves ITD and ILD cues comparatively stable. Because vertical rotations do not substantially alter the left–right spatial relationship between the source and the ears, these binaural cues continue to reflect external source motion rather than a mixture of source and self-motion. Vertical movements do alter the listener’s orientation relative to the source, and may therefore perturb pinna-based spectral cues. Thus, the different precision observed between movement planes may reflect differences in how self-motion perturbs the spatial cues available for estimating source motion.

Consistent with this interpretation, response precision — the trial-to-trial consistency of directional judgements, quantified as the normalised vector-sum length — was significantly higher during vertical than horizontal head movements (M = 0.78 vs. 0.69; paired t-test, t(12) = 3.93, p = .002, dz = 1.09), an advantage shown by 11 of 13 participants. This benefit was specific to precision: directional accuracy (the absolute error of the mean response direction) did not differ reliably between orientations (t(12) = −1.19, p = .256, dz = −0.33) and instead varied with speaker-motion direction (Suppl. Fig. S2). This dissociation is consistent with differential availability or reliability of binaural and spectral cues across the two movement planes; the present design, however, does not isolate these acoustic cues directly, so this remains an interpretation rather than a demonstrated mechanism (see Discussion).

Since participants also moved their heads more slowly in the vertical conditions, one might ask whether the higher precision observed there simply reflects a speed advantage rather than a cue-reliability advantage — consistent with our finding that slower head movements are associated with better compensation. To address this, we repeated the precision analysis controlling for individual trial-level head speed. In this model the vertical-over-horizontal precision advantage remained significant (b = 0.103, 95% CI [0.042, 0.165], t(12) = 3.67, p = .003), whereas within-participant head speed did not predict precision (b = −0.001, p = .61) and did not interact with orientation (p = .32). This indicates that the vertical precision advantage is not explained by movement speed alone and is consistent with a cue-reliability account (Supplementary Fig. S6).

## Discussion

The present study investigated how concurrent head movements influence the perception of moving auditory objects. Across all head-movement conditions, participants showed systematic biases in their perceived direction of sound-source motion, with responses shifted opposite to the direction of head movement. This pattern is consistent with incomplete compensation for self-motion — a failure to fully translate head-centered auditory signals into a stable world-centered percept. Compensation was incomplete in all conditions and was most severely disrupted when the sound source moved along the same axis as the head. In these aligned configurations, participants often reported the source as stationary, suggesting that directional information became highly unreliable when self-motion and source motion are fully confounded, even though participants’ head speed was on average four times higher than the actual speaker speed. Precision was significantly higher during vertical than horizontal head- movement conditions, a pattern consistent with differential vulnerability of binaural and spectral spatial cues to self-motion. Contrary to our prediction, vertical head motion did not produce a reliable overall improvement in directional accuracy; instead, its advantage was specific to response precision.

These findings build on Freeman et al.^10^, who demonstrated systematic compensation deficits during horizontal head movements using a compensation gain framework. In their study, gain values were typically around 0.15 — expressed in the complementary sense used here, this corresponds roughly to an equivalent gain of 0.85 in our study, broadly consistent with our observed range. However, a direct comparison reveals an important distinction: our overall gain estimates reflect an average across all speaker directions, including those where compensation is relatively successful. When speaker motion is aligned with head motion, our data show a collapse of response precision and a loss of reliable directional information in these aligned configurations — rather than a measurable residual gain, since compensation gain is not identifiable when head and source motion are near-collinear (Supplementary Figs. S3 and S5). This is also consistent with the negative relationship we observed between head speed and compensation gain — a result that echoes Freeman et al.’s observation that motion thresholds scale with head speed, implying that one or more of the underlying signals used for compensation becomes less reliable as head speed increases. Whether this unreliability originates from degraded extracochlear signals — such as proprioceptive or vestibular estimates of head velocity — or from the acoustic cues themselves becoming more ambiguous at higher speeds remains an open question. Disentangling these contributions would require manipulations that selectively degrade one signal source while preserving the other, and represents an important direction for future work.

The finding that participants moved their heads far faster than instructed — averaging around 53°/s against a target of approximately 10.5°/s — indicates that participants systematically overshot the instructed target velocity under blindfolded conditions, and may have contributed to undercompensation, although the observed negative association between head speed and gain was exploratory. This difficulty in accurately reproducing a target-matched movement speed is consistent with a broader challenge in coupling self-motion to auditory targets. Leung et al.^11^ observed the complementary error: when actively tracking a moving auditory object, listeners moved their heads substantially slower than the target, and a ramp-up phase was necessary to enable successful tracking at even moderate speeds. Taken together, these findings suggest that the auditory system has poor access to accurate speed information in both directions — whether trying to match or to maintain a prescribed pace — and that this limitation is unlikely to be specific to any one task design. The uncontrolled head speed variability in our data limits the precision of the gain estimates, and a follow-up study with externally paced or mechanically constrained head movements at fixed speeds would provide a cleaner test of both the gain model and the cue-reliability hypothesis.

In most conditions, mean absolute error peaked not exactly at the head-motion direction but slightly off-axis from it — a pattern visible in all conditions except vertical up-to-down. This could reflect a geometric property of the vector-summation model, where near-aligned but not perfectly aligned speaker directions produce the largest angular distortions in the perceived motion direction, or it may indicate that participants retain partial directional sensitivity even for near-aligned motion. Characterizing this directional tuning in greater detail, potentially using finer-grained speaker direction sampling around the head-motion axis, could shed light on the spatial resolution of auditory motion compensation.

A caveat applies to this cue-based interpretation. We attribute the greater precision during vertical head motion to the comparatively stable binaural (interaural time and level difference) cues available when the head rotates in the vertical plane, but we did not manipulate these cues experimentally or derive them from measured or standard head-related transfer functions together with the recorded head and source trajectories. We therefore limit this claim to the statement that the observed pattern is consistent with differential binaural and spectral cue reliability; establishing that these cues mediate the behavioural effect would require an acoustic forward model — measured or standard HRTFs/BRIRs, time-resolved source position and head orientation, predicted ear signals for each trajectory, and derived ITD, ILD and spectral-change measures — linked to behaviour in a participant-aware statistical model.

Taken together, these results reveal that the auditory system attempts to compensate for self- induced motion signals but does so imperfectly, with the degree of failure depending on the axis of head movement, the speed of self-motion, and the geometric relationship between head and source trajectories. The findings parallel well-established results in visual motion perception — such as the Filehne illusion — leading to systematic perceptual biases. That a structurally similar phenomenon exists in audition, despite the auditory system lacking a dedicated peripheral motion sensor analogous to the retina, suggests that motion parsing under self-motion may reflect a general computational challenge shared across sensory modalities.

## Methods

### Participants

Thirteen participants (N = 13, 10 female, mean age 26.15 years) completed the experiment. All reported normal hearing. Each participant completed two experimental sessions of approximately 90 minutes each on two consecutive days. Written informed consent was obtained prior to participation. Experiments were approved by the Ulm University Ethics board.

### Apparatus and Stimuli

Stimuli were presented using a 3 × 3 × 3 m cable robot that is capable of physically moving a loudspeaker (Anker SoundCore Mini) silently at constant velocity in the frontal plane of the participant^12^. Speaker motion directions were equally distributed around a circle, starting at 0° and continuing in 22.5° steps to obtain 16 unique speaker directions. The speed of the speaker was fixed at 33 cm/s for all directions, which corresponds to approximately 10.5 deg/s for a participant with their head at the centre of the workspace. Participants were seated at a distance of approximately 1.8 m from the fronto-parallel plane in which the loudspeaker moved, and the experiment was carried out in a sound-attenuated room.

Each stimulus consisted of a 1.5 s burst of broadband white noise with 250 ms on-/offset ramps, presented at 60 dB SPL measured at the participant’s head position. A stationary loudspeaker situated behind the participant generated a brief, pure tone (500Hz, 800ms, with 200ms on-/offset ramps) to signal the start of each trial and, in the case of head-movement conditions, to prompt participants to begin moving their heads. Following each stimulus presentation, participants indicated the perceived direction of sound-source motion using a joystick ([joystick make and model]). Trials in which the joystick response remained at the neutral position (0,0) were discarded.

Motion tracking markers were placed on the speaker and head of the participants, respectively. We used an OptiTrack motion-capture system (12 infrared cameras) sampling at 120 Hz to continuously track the location and rotation of the speaker and the participant’s head throughout the entire experiment.

### Experiment

The experiment comprised six conditions. In the eyes-open stationary head condition, participants completed 32 trials, consisting of two repetitions for each speaker direction prior to Session 1. Following this, participants were blindfolded and were asked to keep their eyes closed. In the eyes-closed stationary-head condition, participants performed 80 trials per session (160 in total) during which time the loudspeaker was moved 10 times along each of the 16 trajectories. The head-still condition was always administered as the first condition of each session.

The remaining four conditions required participants to move their heads and were run over two days. These head-movement conditions consisted of horizontal left–right, horizontal right– left, vertical up–down, or vertical down–up movements run in a randomized order for each participant. Each of these conditions began with 16 training trials with eyes open, during which participants practiced matching their head speed to the speed of the moving loudspeaker. They were then blindfolded and 160 experimental trials were run (10 repetitions per speaker direction). The order of head-movement conditions was randomized across participants, while making sure that in each day one horizontal and one vertical head movement condition was tested.

At the beginning of each trial, a brief starting tone was presented to cue participants to initiate their head movement. Simultaneously, the loudspeaker began to move, with stimulus playback delayed by 0.73 s to ensure that the stimuli were centered when the sound was played. In this way, the stimulus spanned the period just before and after the speaker crossed the midpoint of the circle. Participants indicated the perceived direction of loudspeaker motion using a joystick. After a response was registered, the loudspeaker returned to a new starting position determined by the next randomly selected movement direction. Because of this randomization, the inter-trial interval varied depending on the length of the repositioning movement.

### Data Analysis

#### Participant’s Head Speed

Head orientation was recorded as quaternions *q*(*t*) and realigned on each trial to a common reference. A reference quaternion *q*_ref *f##*_ was computed from the mean head pose in the initial pre-movement samples and all samples were rotated into this frame via

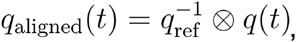

and then converted to Euler angles ϕ(t), θ(t), ψ(t) corresponding to roll, pitch, and yaw, respectively. Angles were unwrapped to avoid discontinuities at ±π.

Only head movements within the duration of stimulus presentation were considered. That is, analyses of head kinematics were restricted to the stimulus window [*t*_on_, *t*_off]_, where *t*_*on*_*= t*_*trial start +*730*ms*_ and *t*_*off*_ *=t*_*on +*1500*ms*_

Angular velocities were obtained by differentiating the unwrapped Euler angles (central differences) and the mean head speed was defined from the net angular displacement in the task-relevant axis divided by the stimulus duration:]

$$\bar{\omega}_{y}=\frac{y(t_{\mathrm{off}})-y(t_{\mathrm{on}})}{t_{\mathrm{off}}- t_{\mathrm{on}}}$$, $$\bar{\omega}_{p}=\frac{p(t_{\mathrm{off}})-

p(t_{\mathrm{on}})}{t_{\mathrm{off}}-t_{\mathrm{on}}}$$,

$$\bar{\omega}_{r}=\frac{r(t_{\mathrm{off}})-r(t_{\mathrm{on}})}{t_{\mathrm{off}}-

t_{\mathrm{on}}}$$ [\deg/\mathrm{ms}].

Head speed was then defined depending on the instructed movement plane:

- Horizontal conditions (hori_left_right / hori_right_left): $$v_{\mathrm{head}} \;=\; 1000\,\bigl|\bar{\omega}_{y}\bigr|\quad [\deg/\mathrm{s}]$$
- Vertical conditions (vert_up_down / vert_down_up): $$v_{\mathrm{head}} \;=\; 1000\,\bigl|\bar{\omega}_{r}\bigr|\quad [\deg/\mathrm{s}]$$
- Stationary_eyes_closed: $$v_{\mathrm{head}}=0$$.

In the head-tracker coordinate frame used here, the up/down sagittal-plane rotation of the vertical conditions registers as roll (ϕ) and the left/right rotation of the horizontal conditions as yaw (ψ); head speed was therefore computed from roll for vertical and yaw for horizontal conditions. An axis audit confirmed that each manipulation was carried overwhelmingly by its intended Euler axis (median |roll| = 46.1°/s for vertical and median |yaw| = 58.8°/s for horizontal motion, each exceeding the next-largest axis by a factor of ∼19–20; Supplementary Fig. S4). We applied three sequential filters to mark and remove trials with inadequate or unreliable head motion during stimulus presentation:

The analysis-window onset was defined during preprocessing as the recorded sound-presentation timestamp plus the configured 730 ms offset. Thin, translucent gray lines show all retained trials from the same participant and horizontal right-to-left movement condition. **(a)** Representative retained trial exhibiting continuous head rotation throughout the analysis window. **(b)** Trial rejected because the total angular change during the initial 500 ms analysis interval (orange shading) was 0.6°, below the required minimum of 10°. **(c)** Trial rejected because it contained 441 ms of consecutively low frame-to-frame angular change (orange shading), exceeding the permitted duration of 350 ms. Low angular change was operationally defined as ≤0.3° per frame, corresponding nominally to an angular speed of ≤36° s^−1^ at the 120 Hz sampling rate. **(d)** Trial rejected as a trajectory outlier because at least one sample fell outside the group mean ±3 SD envelope, calculated from trials remaining after application of the initial-movement and low-angular- change criteria. In panels **a–c**, the highlighted trajectory and each gray reference trajectory were normalized by subtracting their respective head orientation at the analysis-window onset; consequently, these trajectories pass through zero at \(t=0\). In panel **d**, the highlighted trajectory and exclusion envelope were instead referenced to the retained group’s mean orientation at onset, preserving the between-trial orientation offset relevant to the outlier criterion. The gray reference trajectories in panel **d** remain individually onset-normalized. Exclusion criteria were applied sequentially in the order illustrated in panels **b–d**.

##### 1) Insufficient head movement near onset and offset

For each trial, head orientation (roll, pitch, yaw) was examined in two short windows relative to the stimulus: an initial window

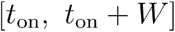

and a terminal window

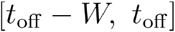

with. Within each window, axis-wise angular excursions were computed as

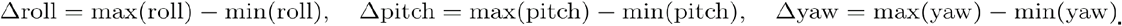

A 3-D movement magnitude was then formed as the Euclidean norm

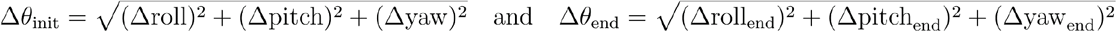

Trials were retained only if both magnitudes exceeded a minimum displacement criterion:

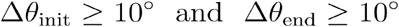

Otherwise, the trial was flagged as “insufficient head movement” and discarded.

##### 2) Prolonged static tracking during the stimulus

To detect tracking dropouts or frozen signals within [*t*_on_,*t*_off_], we computed frame-to-frame changes for both position and rotation:

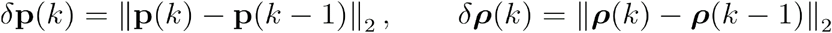

where P= (*x,y,z*) and ρ = (roll,pitch,yaw).

A time sample was marked static if

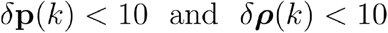

and we computed the duration of each contiguous static run. Trials were removed if any static run during the stimulus window met or exceeded the minimum static-duration threshold (set to 10). Trials without such runs were retained.

##### 3) Time-series outlier rejection relative to the condition mean

For the remaining trials, we constructed a condition-specific head-motion trace during the stimulus window using a single diagnostic angle:

- yaw for horizontal conditions,
- roll for vertical conditions.

Each trace was cropped to a fixed length of 180 samples per trial (1.5s, the stimulus duration). We then computed, across trials, the pointwise mean μ (*t*) and standard deviation. A trial was retained only if its entire trace lay within a ±3σ envelope:

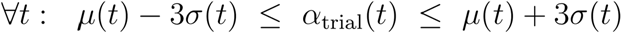

Any violation resulted in removal as a time-series outlier.

Orthogonal to the motion-based filters above, we removed null responses in which the joystick remained at (0,0)(0,0)(0,0) (no perceived motion direction), and any trials with missing/non-finite kinematic samples after alignment and unwrapping.

Filters were applied in the order listed: onset/offset movement, static-tracking detection, time-series outliers. Each stage produced synchronized, trial-level masks applied to both motion and response tables to ensure consistent inclusion across modalities. The number of retained and removed trials at each step was logged.

By applying these filters to the motion data of all participants we rejected 1,988 of 9,346 trials (21.3%): 1,491 for insufficient initial motion, 354 for prolonged static tracking, and 143 as trajectory outliers, leaving 7,358 trials for analysis (per-participant and per-condition counts are given as Supplementary Data; see Supplementary Fig. S4)

### Histogram & Von Mises Fits

The individual responses of all participants for a condition are binned (32 bins, equally spaced around the circle, centered on speaker directions and the angles between speakers). To quantify the perceptual bias in each condition, we fitted a von Mises distribution to the data, which yields parameters μ (mean response direction) and κ (analogous to 1/σ^2^), similarly to a normal distribution. A κ value of 0 indicates a uniform distribution, whereas larger κ values reflect a stronger concentration of responses around μ. von Mises distributions are fitted to the raw response angle data.

### Gain Factor

For each experimental condition, trials were grouped by the speaker motion direction. Within each group, we fit a von Mises distribution to the participants’ response angles (given by joystick directions) and took the fitted location parameter as the empirical mean response direction. That is, for each speaker motion direction we fit a von Mises distribution using all participants’ responses for this speaker direction:

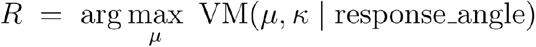

using *scipy*.*stats*.*vonmises*.*fit()* followed by wrapping to [0,2π). We use this averaged value of responses to one speaker direction to achieve a more stable and reliable response direction, leading to more stable gain calculations.

We then modeled speaker and head motions as 2-D motion vectors with magnitudes given by their angular speeds:

- Speaker motion vector: **S** = *s* · [cos *W*, sin *W*], where *s* is the speed of the speaker, fixed to 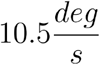
- Head motion vector (per group, averaged across the grouped trials):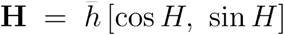, where *H* is the head motion direction in the given condition and 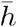 the mean calculated angular velocity of the head over the grouped trials.

The speaker motion vector represents the true motion of the speaker in the world, i.e., the world-centered velocity of the speaker. Subtracting the head-motion vector from the world- centered source-motion vector provides a kinematic approximation of source motion in head- centered coordinates. Since this varies from the response of participants, we assumed that perceived motion results from vectorial combination of world-centered source motion and a residual head-motion component scaled by a compensation gain *g* .

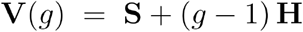

That is, *g =* 1 corresponds to perfect subtraction of head motion (full compensation), *g =* 0 to no compensation (head motion fully contaminates perception), and *g* > 1 to over- compensation.

## Gain Factor

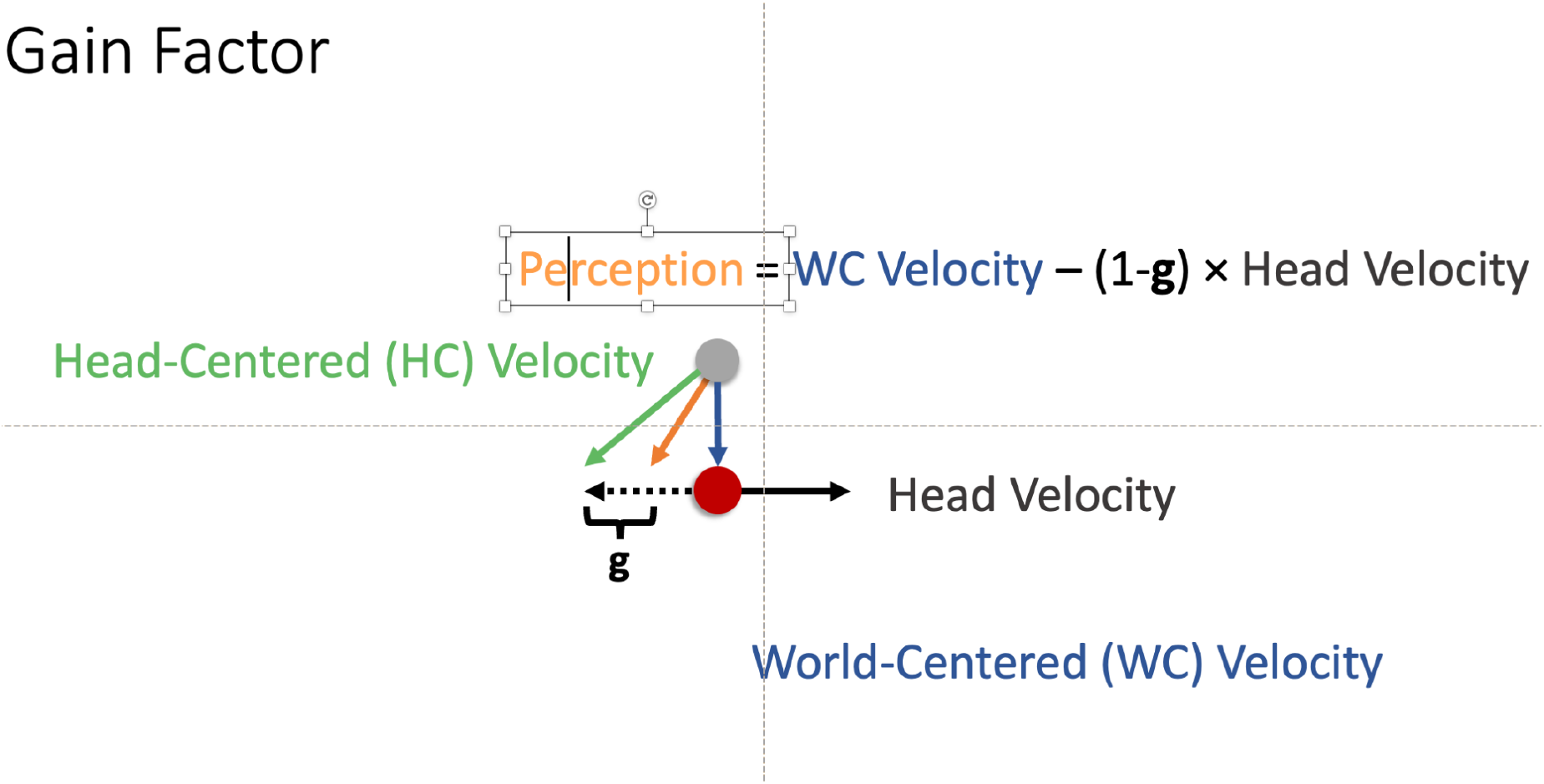

Fig. Gain factor calculations

By transforming this motion vector *V (g)* to a response angle by atan2 we obtain an estimated response 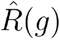:

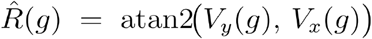

We then estimated the best fitting *g* for given *R,H* and *S*. This is, for each (condition,W)(\text{condition}, W) group, we estimated by solving for zero wrapped angular discrepancy between the model prediction and the empirical mean response:

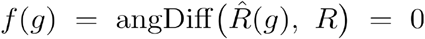

This scalar root was obtained with Brent’s method (*scipy*.*optimize*.*root_scalar()*). For a group of responses (for one speaker direction) we get the same *g* values.

### Handling undefined configurations

Head–source collinearity provides no information to separate from. Accordingly, after gain estimation we removed trials if the speaker direction matched the head-motion direction or its opposite:

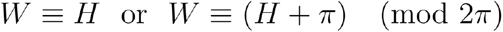

These trials are excluded from summary statistics of gain (mean, s.d., min, max) reported per condition.

For each condition we report the distribution of compensation gains across the 16 speaker directions, yielding a single per speaker direction and condition, grounded in the von-Mises-based empirical direction and a physically interpretable vector model of source and head motion.

All angles were expressed in radians. We used canonical wrapping to [0,2π), wrap_to_2π (α) α mod 2π and smallest signed angular differences in (−π,π], angDiff (a,b) = (*a* − *b* + π) mod 2π − π to ensure numerically stable comparisons between circular quantities.

To characterize directional variation in compensation strength, we performed an additional exploratory gain analysis in which gain was estimated separately for each speaker direction. Using the processed trial-wise dataset from all non-stationary conditions, we pooled trials either across all motion conditions, across horizontal and vertical conditions separately, or within each condition. For a given speaker direction, we fit a single scalar gain parameter by minimizing the mean squared circular error between the observed response angle and the angle predicted by a vector-summation model combining speaker motion and head motion. The speaker-motion vector had fixed direction and nominal speaker speed, whereas the head-motion vector used the condition-specific head-motion direction and the trial’s mean angular head speed. Gain estimation was performed with bounded scalar optimization. To summarize variability across participants, we repeated the fit separately for each participant within each speaker-direction subset and report the participant-level median gain together with the interquartile range. In additional control analyses, trials in which speaker motion and head-motion direction were aligned were optionally excluded to assess whether such geometries disproportionately influenced extreme gain estimates.

### Statistical analysis

Unless stated otherwise, all analyses were performed in Python 3.13 using NumPy, SciPy, pandas, and statsmodels. Circular quantities (response directions, biases, and gains) were treated with the wrapping and angular-difference conventions defined above, and all group- level statistics were computed across participants (n = 13).

Response accuracy was quantified as the mean absolute error (MAE), defined as the angular difference between the reported and the true speaker-motion direction and averaged across participants for each speaker direction; the reported error bars denote 95% confidence intervals (1.96 × standard error across participants). Response precision was quantified as the normalised vector-sum length (range 0–1) computed per participant and condition.

To compare precision between horizontal (Hori L-R and Hori R-L combined) and vertical (Vert U-D and Vert D-U combined) head-movement conditions, we used a two-tailed paired- samples t-test on participant-level mean precision. To test whether this difference reflected cue reliability rather than the slower head speeds observed in the vertical conditions, we repeated the comparison while controlling for trial-level head speed (a participant-level linear model with participant and speaker-direction fixed effects and participant-clustered standard errors, head speed centred within participant; Supplementary Fig. S6).

The relationship between head speed and compensation gain was assessed with a Pearson correlation, using gain estimates restricted to trials with speaker motion perpendicular to the head-motion axis and binned into 10 equal-sample head-speed bins (n = 632 trials per bin). Statistical significance was assessed at α = 0.05.

