## Supplementary material for "Moving Heads, Moving Sounds: Systematic Errors in Auditory Motion Perception": Supplemantary

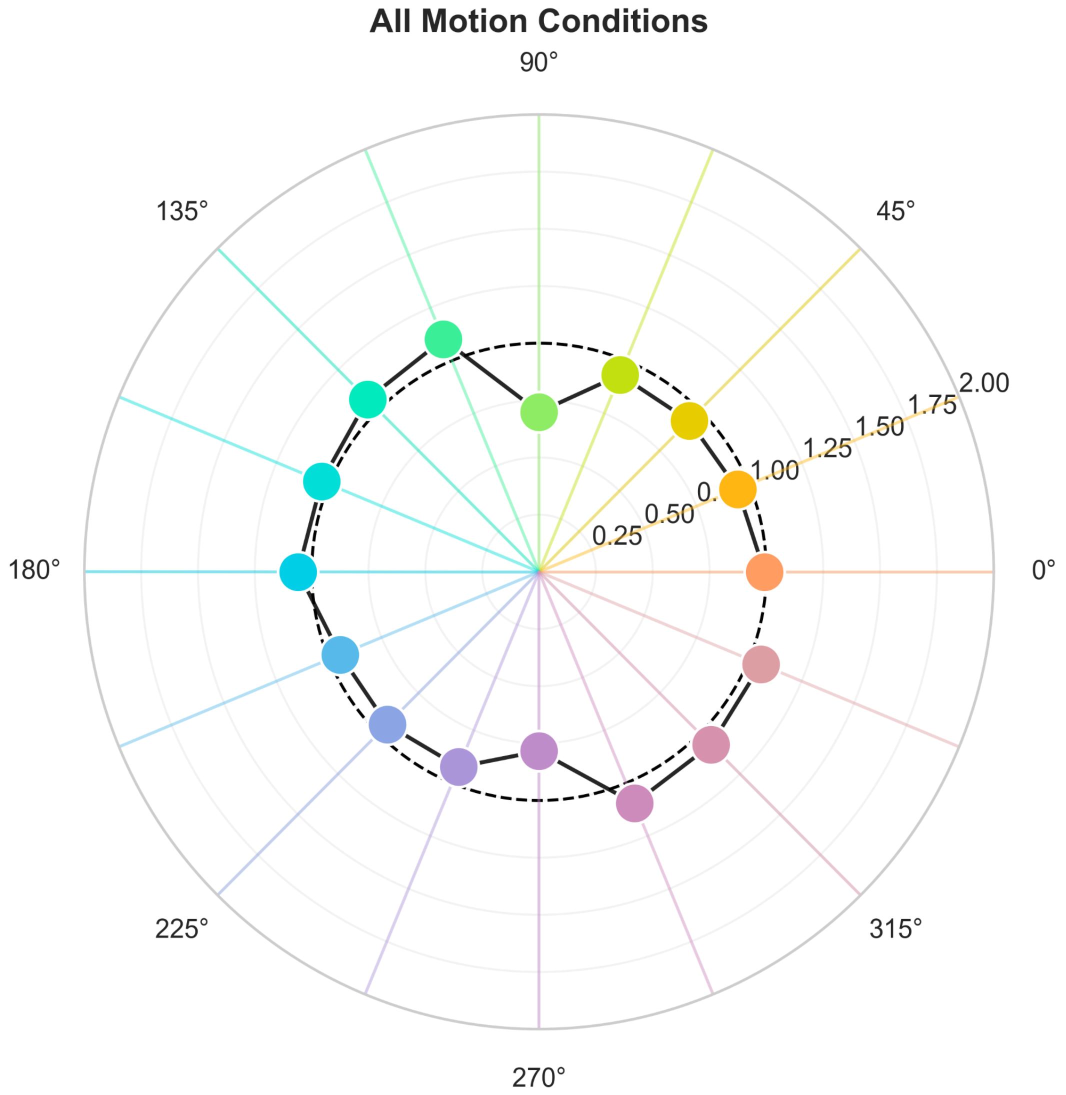

**Supplementary Figure S1.** These per-direction gains are pooled across participants and conditions and therefore differ from the condition-level estimates in the main text and the participant-level estimates in Supplementary Fig. S3; this pooled summary is descriptive only and is not used for statistical inference.

Polar summary of experimentally estimated gain as a function of speaker direction, pooled across all motion conditions (hori_left_right, hori_right_left, vert_up_down, vert_down_up). Each angular position corresponds to one speaker direction in the response-centered coordinate system, and each colored dot indicates the participant-level median fitted gain for that direction after pooling trials across conditions. The black radial bar shows the interquartile range across participant-level gain estimates (Q1 to Q3), providing a robust measure of between-participant variability. The dashed circle marks unity gain (gain = 1), that is, the prediction of a veridical compensation model. Values above 1 indicate overcompensation, whereas values below 1 indicate undercompensation. The black line connects medians across neighboring speaker directions to emphasize directional trends, but does not imply a continuous underlying function. Colored radial guide lines indicate the sampled speaker directions. Gains were estimated by fitting a single scalar gain parameter that minimized circular localization error between observed responses and the vector combination of speaker motion and head motion, using each trial's mean head angular speed. Because the fit is performed separately for each speaker direction after pooling across motion conditions, the figure captures directional structure in gain independently of condition-specific summaries. Large spreads for some directions indicate substantial between-participant heterogeneity and/or reduced parameter stability for those geometries, so these values should be interpreted as descriptive rather than as evidence for a smooth directional tuning function.

### Dissociating response precision from directional accuracy

Precision and directional accuracy describe distinct aspects of performance. Precision (response consistency) quantifies how reproducible repeated responses are, irrespective of whether their mean is centred on the true speaker-motion direction, whereas accuracy quantifies how close the mean response direction lies to the target. A condition can therefore yield highly consistent responses that are systematically displaced from the target — high precision but low accuracy. To clarify the interpretation of the vector-sum analyses, we examined the two measures separately.

For each of the 13 participants, four head-movement conditions, and all 16 speaker-motion directions (0–337.5° in 22.5° steps; the stationary condition was excluded), precision was quantified as the normalised vector-sum length of the reported directions (0–1, higher = more consistent) and accuracy as the absolute circular difference between the participant's mean response direction and the presented speaker-motion direction (lower = more accurate). The two opposing conditions within each orientation were averaged (vertical: up-to-down and down-to-up; horizontal: left-to-right and right-to-left), and the 16 directions were then averaged with equal weight to give one vertical and one horizontal value per participant. Orientations were compared with two-sided paired-samples t-tests, with effect sizes reported as Cohen's dz.

Precision was higher during vertical (M = 0.776, SD = 0.088) than horizontal (M = 0.688, SD = 0.129) head motion; the mean paired difference was 0.088, 95% CI [0.039, 0.137], t(12) =

3.93, $p = .002$, $d_z = 1.09$. Eleven of 13 participants, and 12 of 16 speaker directions, showed higher vertical precision.

Directional accuracy did not differ reliably between orientations (vertical $M = 39.40°$, $SD = 8.11°$; horizontal $M = 42.88°$, $SD = 14.05°$; difference −3.49°, 95% CI [−9.85°, 2.88°], $t(12) = −1.19$, $p = .256$, $d_z = −0.33$). Vertical motion produced lower mean error for 10 of 16 directions, but the orientation curves crossed repeatedly across the direction range, indicating that accuracy depends on the specific combination of head-motion orientation and speaker-motion direction rather than on orientation alone.

This dissociation also explains an apparent discrepancy with the pooled polar vector-sum plot (Fig. 1D), in which several horizontal mean-response directions align closely with their target spokes and can create an impression of better horizontal accuracy. In the polar representation the angular position of each point encodes the mean response direction (and its distance from the target spoke, the directional error), whereas the radius encodes precision; horizontal points tend to have shorter radii, i.e. lower precision. The polar plot additionally pools trials across participants before computing each vector sum, which does not preserve participants as independent units, so the participant-level paired analysis reported here is the more appropriate basis for inference. The black diamond in that figure should likewise not be read as an overall precision estimate, as its radius reflects the concentration of the 16 mean-response angles rather than trial-to-trial response precision.

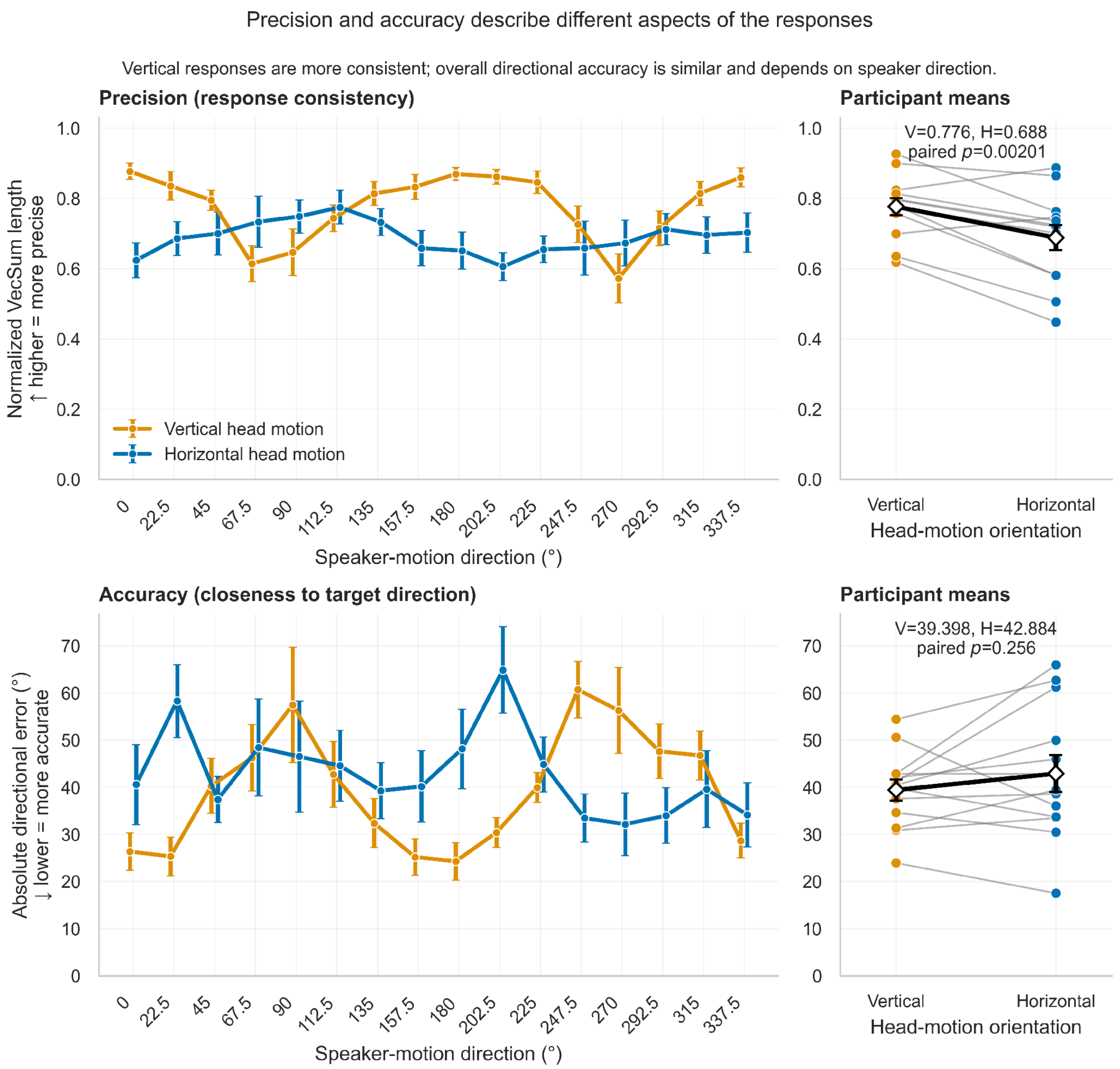


**Supplementary Figure S2.** Precision and directional accuracy for vertical and horizontal head motion across all speaker-motion directions. Upper panels: response precision (normalised vector-sum length; higher = more consistent). Lower panels: absolute directional error (lower = more accurate). Left panels show participant-level group means ± SEM for each of the 16 speaker-motion directions; right panels show participant averages across all directions, with grey lines connecting the same participant, coloured circles for individual participant means, and black diamonds ± SEM for the group mean. Precision was significantly higher during vertical than horizontal head motion ($t(12) = 3.93$, $p = .002$, $dz = 1.09$), whereas absolute directional error did not differ significantly between orientations ($t(12) = -1.19$, $p = .256$, $dz = -0.33$). Response precision and directional accuracy are thus separable: vertical head motion increased response consistency without a corresponding overall improvement in accuracy.

### Participant-level compensation gain and geometric identifiability

The compensation gain reported in the main text (Fig. 2) was estimated from responses pooled across participants and is poorly identified for head/source geometries that approach collinearity. Because identifiability degrades continuously as the speaker-motion and head-motion vectors approach alignment — rather than failing only at exact collinearity — we refit the model $V(g) = S + (g - 1)H$ with one gain per participant and movement condition, estimated directly from the trial-wise circular responses by minimising the mean squared circular prediction error, so that the participant is the unit of inference. Geometric identifiability was quantified as $|\sin(\theta\_speaker - \theta\_head)|$, and gains were examined across a range of minimum-identifiability thresholds, with a primary threshold of 0.5 (a minimum separation of 30° between head- and source-motion directions).

At the primary threshold, mean participant-level gains were 0.89 (horizontal left-to-right), 0.79 (horizontal right-to-left), 0.88 (vertical up-to-down) and 0.91 (vertical down-to-up), confirming systematic undercompensation (gain < 1) in every condition and closely matching the pooled estimates in the main text. Gains were stable across identifiability thresholds except for near-collinear geometries, where estimates became extreme and reached the optimisation bounds — behaviour expected from the loss of identifiability in that geometry rather than from a change in perception. Aligned and opposite head/source configurations are therefore not interpreted through gain; the corresponding perceptual breakdown is described using precision and directional-error measures instead.

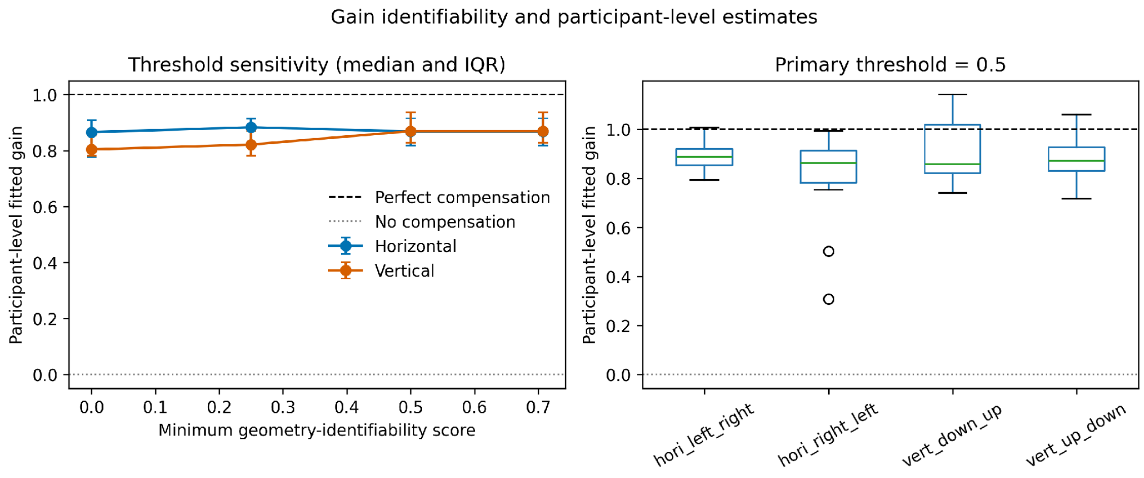


**Supplementary Figure S3.** Participant-level compensation gain and its geometric identifiability. Gain was fitted separately for each participant and movement condition from trial-wise circular responses. Identifiability is quantified as $|\sin(\theta\_speaker - \theta\_head)|$; panels show gain estimates and their stability as a function of the minimum identifiability threshold applied, with the primary threshold of 0.5 (≥30° head–source separation) marked. Mean gains remain below unity across conditions and are stable except near collinear geometry, where estimates become unidentifiable.

### Head-motion axis audit and trial-exclusion accounting

To verify the coordinate convention used for head speed, we measured which tracker Euler axis carried each manipulation. Vertical conditions were dominated by roll (median |roll| = 46.1°/s, versus |pitch| = 2.2°/s and |yaw| = 2.3°/s), and horizontal conditions by yaw (median |yaw| = 58.8°/s, versus |roll| = 2.9°/s and |pitch| = 3.1°/s); in each case the intended axis exceeded the next-largest axis by a factor of approximately 19–20. Head speed was therefore computed from roll for vertical and from yaw for horizontal conditions.

The three sequential motion filters removed 1,988 of 9,346 trials: 1,491 for insufficient initial motion, 354 for prolonged static tracking, and 143 as trajectory outliers, leaving 7,358 trials for analysis. Per-participant and per-condition counts are provided as Supplementary Data; exclusions were concentrated in a small number of participants who struggled to sustain the instructed movement rather than being spread evenly, while most participants retained the large majority of their trials. The vertical-over-horizontal precision advantage was present both before and after the trajectory-outlier filter (vertical vs. horizontal precision: 0.74 vs. 0.66 using all trials; 0.78 vs. 0.69 using retained trials), indicating that the effect is not an artefact of the exclusion procedure.

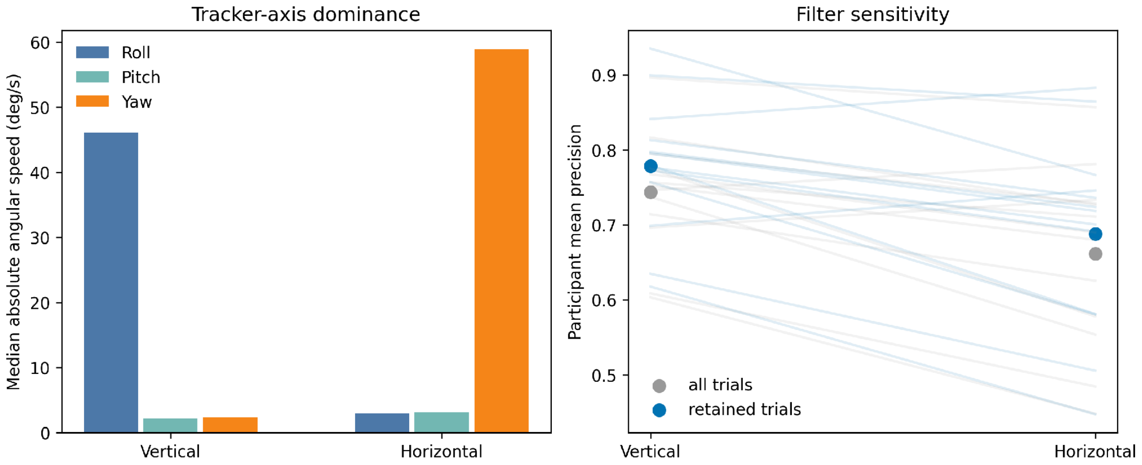


**Supplementary Figure S4.** Head-motion axis audit and filter sensitivity. Left: median absolute angular speed in each tracker Euler axis (roll, pitch, yaw) for vertical and horizontal conditions, showing that each manipulation is carried overwhelmingly by its intended axis (vertical = roll, horizontal = yaw). Right: participant mean precision (normalised vector-sum length) for vertical and horizontal head motion computed on all trials and on retained trials only, showing that the vertical precision advantage is preserved regardless of the trajectory-outlier filter. Full per-stage and per-participant exclusion counts are reported in the text and provided as Supplementary Data.

### Neutral (null) joystick responses

Trials in which the joystick remained at the origin were excluded from the directional analyses because the reported angle is numerically unstable near (0,0). Because participants reported that the source sometimes appeared stationary when head and source motion were aligned, we additionally examined neutral responses as an outcome in their own right. Exact neutral responses were rare: 10 of 8,306 movement trials, occurring in 3 of 13 participants, with no systematic concentration in aligned geometries (the highest rate was 4 of 130 trials in the vertical down-to-up aligned cell). Near-neutral responses (joystick radius ≤ 0.01–0.1) are reported as sensitivity analyses. Because no response dead-zone or explicit "no-motion" option was defined a priori, these events cannot be interpreted as perceived-stationary reports, and they are far too infrequent to account for the precision collapse observed in aligned conditions; that collapse instead appears as broad, systematically biased response distributions.

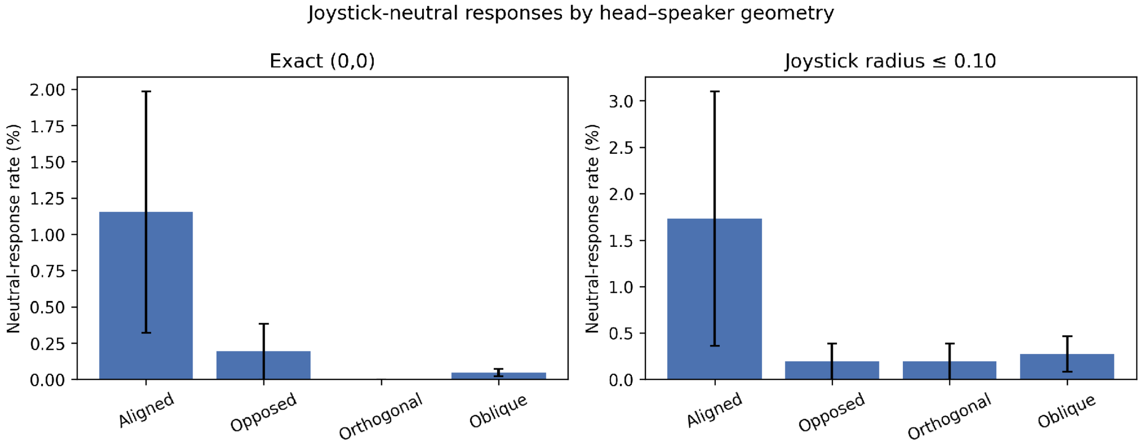


**Supplementary Figure S5.** Neutral joystick responses as an outcome. Rate of exact and near-neutral joystick responses by head/source geometry (aligned, oblique, opposed, orthogonal) and movement condition, shown for exact neutral responses and for a near-neutral threshold (joystick radius ≤ 0.10). Neutral responses are rare and not concentrated in aligned geometries, and are therefore not treated as perceived-stationarity reports.

### Head speed and precision: participant-level model

To test whether the greater precision during vertical head motion reflects cue reliability rather than the slower head movements observed in vertical conditions, we modelled precision (normalised vector-sum length per participant × condition × speaker direction) with participant and speaker-direction fixed effects and participant-clustered standard errors, including within-participant-centred head speed, head-motion orientation, and their interaction; cells with fewer than three valid trials were excluded. The vertical-over-horizontal advantage was significant ($b = 0.103$, 95% CI [0.042, 0.165], $t(12) = 3.67$, $p = .003$). Within-participant head speed did not predict precision ($b = -0.001$, $p = .61$), and the speed × orientation interaction was not significant (p = .32). Participant-level slopes of precision on head speed scattered around zero in both orientations. The precision advantage for vertical head motion is therefore not explained by differences in head speed.

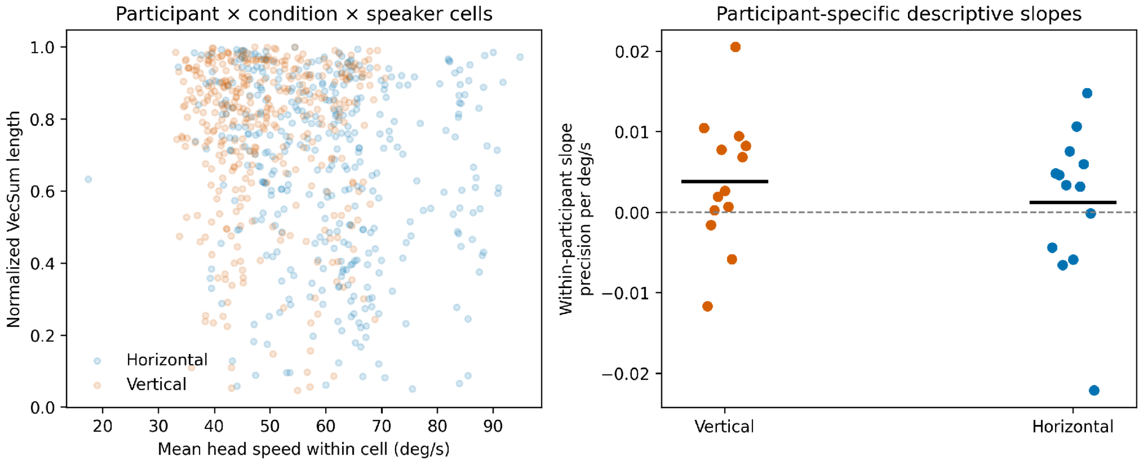


**Supplementary Figure S6.** Head speed and precision at the participant level. Precision as a function of within-participant head speed for vertical and horizontal head motion, with participant-level regression slopes and the fitted orientation effect from a participant-fixed-effect model with participant-clustered standard errors. The vertical precision advantage persists after accounting for head speed, and head speed itself does not reliably predict precision within participants.
